# Genetic evidence for parthenogenesis in small carpenter bee, *Ceratina dallatoreana* in its native distribution area

**DOI:** 10.1101/2022.05.30.494075

**Authors:** Michael Mikát, Jakub Straka

## Abstract

Arrhenotoky is typical mode of reproduction for Hymenoptera – females originate from fertilized eggs, males from unfertilized eggs. However, some lineages of Hymenoptera have switched to thelytoky, where diploid females originate instead from unfertilized diploid eggs. In the contras with some other hymenopteran lineages, thelytoky is generally very rare in bees.

Here, we examined reproduction in the small carpenter bee *Ceratina dallatoreana*, which is assumed to be thelytokous. We compared genotype of microsatellite loci between mothers and their offspring. Offspring were genetically identical with mother in all cases. We did not detect any male offspring. Therefore, we conclude that parthenogeny is the prevailing, and perhaps obligate, mode of reproduction in *C. dallatoreana*. Offspring were clones of their mother with no observed decrease of heterozygosity. Thus the cytological mode of parthenogenesis is apomixis, or automimic with central fusion and extremely reduced or non-existing recombination.

*Ceratina* bees are originally facultatively eusocial, therefore thelytoky may influence social evolution by causing extremely high within-colony relatedness. However, to date no multifemale nests have been recorded in *C. dallatoreana*.

## Introduction

Sexual reproduction predominates in animals, however, parthenogenesis has evolved repeatedly in many lineages (Engelstädter, 2008; Gokhman & Kuznetsova, 2018; Neiman & Schwander, 2011; Normark, 2003; Thierry, 2013). Strictly parthenogenetic lineages are usually young (Fujita et al., 2020; Neiman et al., 2009), and though obligate parthenogenesis can be successful in short-term, in the long-term sexual reproduction is more successful (Neiman & Schwander, 2011; Thierry, 2013). Parthenogenesis probably most often evolved not as an adaptive trait, but as by a product of hybridization or manipulation by a symbiont (Tvedte et al., 2019; Vavre et al., 2004). In insects the occurrence of parthenogenesis differs between orders, and is especially common in stick insects and mayflies (Liegeois et al., 2021; Tvedte et al., 2019).

Parthenogenesis is very diverse and different types of parthenogenesis most likely evolved by different evolutionary mechanisms, each having different consequences for the genetics of the taxa in which it evolved (Engelstädter, 2008). In obligate parthenogenetic population only females exist. However, in many species parthenogenesis is conditionally dependent and present together with sexual reproduction in the same population (Liegeois et al., 2021; Normark, 2003). Such facultative or cyclical parthenogenesis is more common than obligate parthenogenesis (Hörandl et al., 2020).

There exists several cytological mechanisms of parthenogenesis, which strongly differ in influence on genetic diversity and heterozygosity of offspring (Engelstädter, 2017; Hörandl et al., 2020; Pearcy et al., 2006). Parthenogenesis can be caused by absence of meiosis (mitotic parthenogeny, apomixis) or with meiosis, but diploidy restored by various mechanisms (meiotic parthenogenesis) (Stenberg & Saura, 2009). Mitotic parthenogenesis causes increased heterozygosity, since recombination leading to loss of alleles does not occur (Schwander & Crespi, 2009; Tsutsui et al., 2014; Tvedte et al., 2019). On the other hand, meiotic parthenogeny should lead to decreased heterozygosity, because diploidy is restored by endomitriosis (leading to zero heterozygosity) or fusion of products of meiosis, having a similar outcome to self-fertilization – heterozygosity of offspring may be the same or smaller than that of a parent in each locus, and it is always lower on average. Typically, automictic parthenogenesis with terminal fusion (fusion of sister pronuclei) leads to a rapid decrease of heterozygosity (Alavi et al., 2018; Engelstädter, 2017). However, heterozygosity can be retained in the case of meiotic parthenogeny with central fusion, if crossing over is not present in meiosis (Engelstädter, 2017; Stenberg & Saura, 2009). In these cases, there always merge different products of meiosis I, which contain complementary half of genetic information of mother. Result of meiotic parthenogeny with central fusion without recombination is clone of mother similarly as in mitotic parthenogeny (Engelstädter, 2017).

For Hymenoptera, a haplodiploid sex determination system (Kooi et al., 2017; Normark, 2003). Male offspring originate from unfertilized eggs, and are therefore haploid, females from fertilized eggs, and are therefore is diploid (Kooi et al., 2017; Stubblefield & Seger, 1994). Unmated females thus produce only male offspring (Shukla et al., 2013) and the sex of offspring depends the decision to fertilize an egg or not in mated females (Gerber & Klostermeyer, 1970; Stubblefield & Seger, 1994). Arrhenotoky can be a predisposition for evolution of thelytoky (Vorburger, 2014). Thelytokous reproduction has evolved repeatedly in many hymenopteran lineages (Kooi et al., 2017; Vorburger, 2014). It is well-documented in many sapflies and parasitic Hymenoptera (especially Chalcidoidea, Cynipoidea and Ichnumoidea), but is found less frequently within aculeate Hymenoptera (Kooi et al., 2017). Within aculeate Hymenoptera, the best evidence tor thelytoky is from social species (Goudie & Oldroyd, 2018; Wenseleers & Van Oystaeyen, 2011). Thelytoky evolved repeatedly in ants and has been documented in at least 50 species to date (Goudie & Oldroyd, 2018; Heinze, 2008; Rabeling & Kronauer, 2013). In bees, facultative thelytoky is known in *Apis mellifera capensis*, where workers lay thelytokous eggs, usually not in the nest that they originated from (Goudie & Oldroyd, 2014, 2018). However, thelytoky is very infrequent in solitary nesting and weakly social Hymenoptera, with few exceptions (Kooi et al., 2017). Based on sex ratio, thelytoky is assumed in several *Ceratina* species, including *C. dallatoreana* (Daly, 1966; Snelling, 2003). Parthenogeny was considered also for several Australian Halicitd bees, however probably these species are normally sexually reproducing and evidence for parthenogeny is insufficient (Michener, 1960).

*Ceratina dallatoreana* (Apidae: Xylocopinae) nests in broken dead stems with pith (Daly, 1966). Within these stems females build a series of linear cells (Daly, 1966). Although facultative sociality is common in this genus (Groom & Rehan, 2018; Rehan et al., 2009; Sakagami & Maeta, 1977), social nests have not been detected in this species to date (Daly, 1966; Mikát et al., 2022). This species is distributed in the most of Mediterranean and also in central Asia (Terzo, 1998; Terzo and Rasmont, 2004, fig 1). It also exists as an introduced species in California (Daly, 1966). Males are extremely rare in this species (Daly, 1966, 1983), therefore it is assumed that *C. dallatoreana* reproduce by thelytokous parthenogeny.

**Fig 1:**
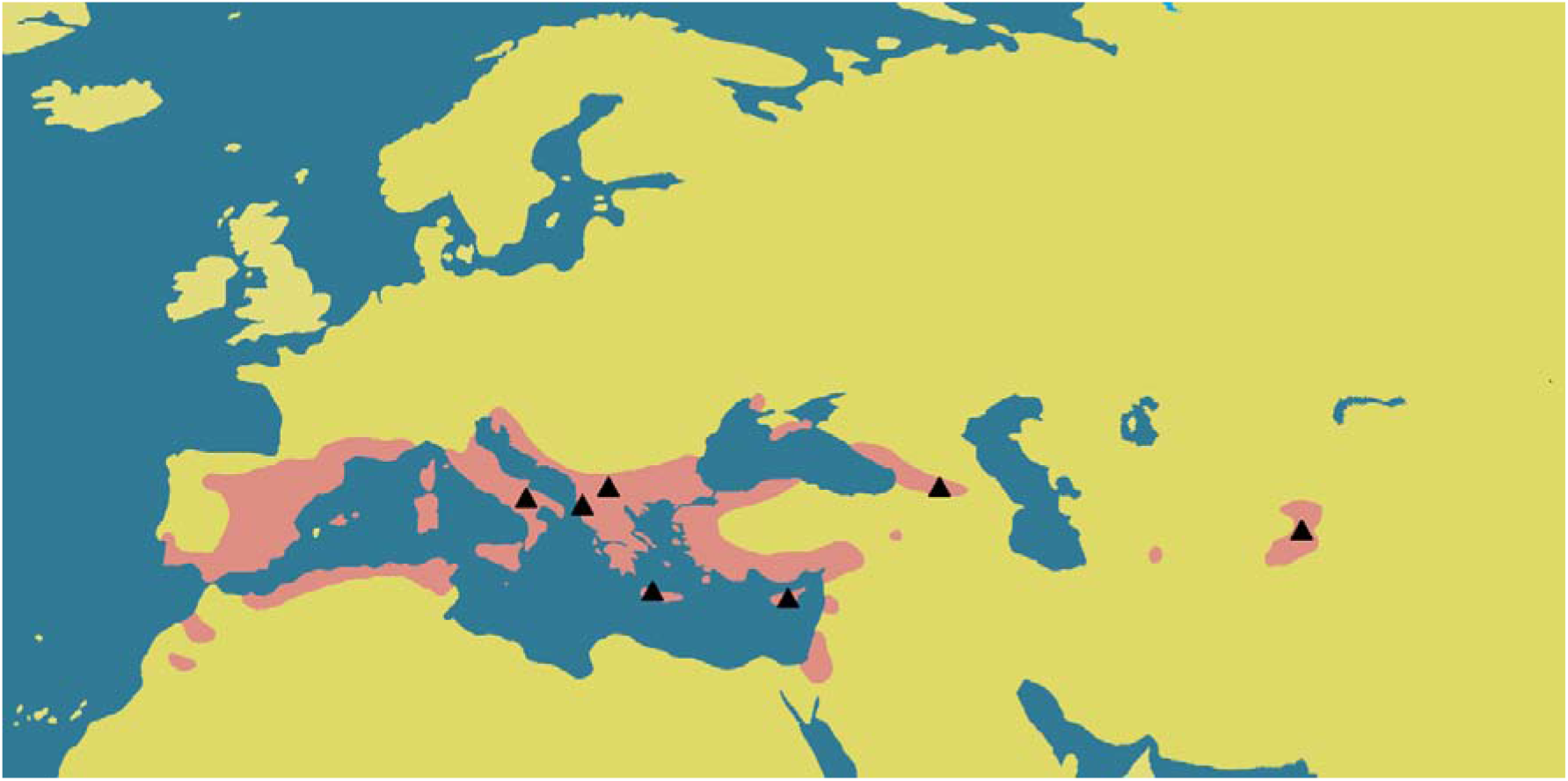
Native range of *Ceratina dallatoreana*. Map is based on (Terzo & Rasmont, 2004, 2011) and new localities from this study. Red – range of *C. dallatoreana*, yellow – other lands, blue – sea and large lakes, Black triangles – sites where *C. dallatoreana* samples for this study were collected.

In this study, we use microsatellite genetic markers for examination of presence and density of parthenogenesis in different populations across the known range of *C. dallatoreana* and assess if heterozygosity is in Hardy-Weinberg equilibrium, increased or decreased in this species.

## Methods

We collected nests of *C. dallatoreana* in several locations across its area of distribution. Nests were collected in Cyprus (2018 and 2019), Italy (regions Puglia and Lazio, 2013 and 2017), Greece (Crete Island, 2018 and 2020), Albania (2018) and Tajikistan (2019). Coordinates of locations, where nests were sampled are shown in supplementary dataset. Additionally, we analyzed individuals collected in Georgia (2013-2014) and Northern Macedonia (2014)

Nests were collected from natural nesting opportunities, in stems which were broken or cut by human management. In Cyprus and Crete we cut some stems to increase nest density for ease of sampling. The most common nesting substrate were *Rubus* spp. and *Foeniculum vulgare*, however nests in dead stems of other plants nests were also collected. Nests were collected in evening (after 18:00 of local time) to ensure that all inhabitants were inside the nest. Nests were open by clippers, and nest contents (number of adults and number and stage of juveniles) were noted. Individuals were preserved in 96% ethanol for further analysis.

For analysis of relatedness we used only nests in active brood nest and full brood nest stages. Active brood nests contain currently provisioned brood cells with a pollen ball, with or without an egg, in the outermost brood cell (Mikát et al., 2021; Rehan & Richards, 2010). Full brood nests contain larva or pupa in innermost and outermost brood cell, as females have already completed provisioning, and are now guarding their offspring until adulthood (Mikát et al., 2021; Rehan & Richards, 2010). We used these two stages, because for these stages it is unambigious that exchange of individuals between nests did not occur.

In total, 59 nests with 188 offspring were analyzed, with 2 nests and 10 offspring from Albania, 6 nests and 23 offspring from Crete, 32 nests and 89 offspring from Cyprus, 10 nests and 38 offspring from Italy and 9 nests and 28 offspring from Tajikistan.

### Isolation of DNA

We isolated DNA using Chelex protocol. Isolation was done usually from part of an individual (one or two legs from adults or pupae, part of the body of most larvae), but whole eggs and whole bodies of small larvae were also used. Samples were transferred to microcentrifuge tubes and dried for at least three hours. Later, we added 8 μl of proteinase K and 50 μl of 10% Chelex solution. This mixture was vortexed and inserted into a thermo cycler. The mixture was heated to 55°C per 50 mins and 97°C per 8 mins then cooled. The whole mixture was then vortexed and inserted into a centrifuge. After this 30 μl of supernatant transferred to a well in the PCR plate.

### Optimalization of multiplex

We selected 12 female *C. dallatoreana* (9 from Cyprus and 3 from Tajikistan) for testing of microsatellite loci. We used microsatellite primers developed for *C. nigrolabiata* (Mikát et al., 2019). Fourteen microsatellite loci were arranged in two multiplexes. The first multiplex was previously applied to *C. nigrolabiata, C. chalybea* and *C. cyanea* (Mikát et al., 2019). The second multiplex contained six loci. Four were different than loci in first multiplex (17 and 36 marked by 6FAM, 9 marked by VIC and 7 marked by PET), and two loci (12 and 51) were shared with the first multiplex but marked by a different color.

We evaluated results of amplification and obtained four possibilities for each locus: a) a locus was successfully amplified in all cases and was polymorphic b) a locus was successfully amplified in almost all cases and was polymorphic c) a locus was amplified in all cases, but was not polymorphic, d) amplification of locus failed (this occurred in a high proportion of cases, table S1).

Polymorphic and reliable loci were 30, 23, 8, 67,17,36,9,12 (table S1, number system of loci is same as for *C. nigrolabiata*, (Mikát et al., 2019)). However, we excluded locus 30 for overlap with same color marked loci and locus 8 for interaction of primers with primers for another locus. Six microsatellite loci were thus retained for final analysis (Table S1, S2).

### PCR and Fragmentation analysis

We used Type-it Multiplex PCR Master Mix (Quiagen) according the manufacture’s protocol. Primers of six microsatellite loci were use in concentration of 0.05 μmol/l. We used these PCR conditions: 95°C for 15 minutes; 30 cycles of 94°C for 30 s, 60°C for 90 s, 72°C for 60 s; and finally, 60°C for 30 min. After PCR, we mixed 0.8 μl of PCR product with 8.8 μl of formamide and 0.4 μl of marker Liz 500 Size scanner (Applied Biosystems). We heated mixture to 95°C for 5 min and then cooled to 12°C. Fragmentation analysis was performed on a 16-capillary sequencer at the Laboratory of DNA Sequencing at the Biological section of Faculty of Science, Charles University, Prague. Identification of alleles was performed in Gene Marker (Soft Genetics) software.

### Analysis of ploidy and heterozygosity

We included mothers from nests and addition individuals in this analysis. We did not include offspring (as they had same genotypes as mothers) to this analysis. For each locus, we checked if an individual had one allele (homozygote) or two alleles (heterozygote). Individuals were considered to be diploid when heterozygous in at least one locus. Individuals which had only one allele in each locus were considered to be haploid. We analyzed 132 females (30 from Crete, 64 from Cyprus, 11 from Georgia, 12 from Italy, 9 from Tajikistan, 3 from Albania and 3 from Northern Macedonia). We also analyzed one gynandromorph (individual with female morphology of head and male morphology of abdomen) from Tajikistan.

### Analysis of deficit or surplus of heterozygotes

We used mothers and additionally sampled females for this analysis. We tested females from two populations, Lefkara village, Cyprus (N=40), and Georgiopoli village, Crete (N=26). All individuals were collected maximally ten kilometers from each other in the same population We calculated observed and expected heterozygosity using software Genepop, version 4.7.5. (Rousset, 2020). Finally, we tested the possible excess or deficiency of heterozygotes also using Genopop.

### Analysis of parthenogeny

We compared the genotype of each mother with the genotypes of their offspring from the same nest, checking if they shared the same genotype. We analyzed 188 offspring from 58 nests in total. Ten offspring from two nests were from Albania, 23 offspring from six nests were from Crete, 89 offspring from 32 nests from Cyprus, and 28 offspring from 9 nests form Tajikistan.

## Results

### Ploidy

All analyzed females from maternal generation (n = 136) were heterozygotes in at least one loci. One female was a heterozygote in only one locus, other females were heterozygotes in at least two loci. Thus we determined that *C. dallatoreana* females are diploid.

In Tajikistan, we collected one gynandromorph individual. This gynandromorph was homozygote in all loci, therefore we considered this individual as haploid.

### Heterozygosity

We detected generally high heterozygosity in our studied loci. Average heterozygosity across all locations and loci was 56.25%. However, heterozygosity differs between loci, with the highest in locus 36 (97.06%), and lowest in locus 12 (4.41%, Table 1). Proportion of each locus across different geographical areas is shown in Table 1.

**Table 1:** Proportion of heterozygotes in each studied loci by geographical area. Other includes samples from Albania (N=3) and Northern Macedonia (N=3).

| Country | N | Proportion of heterozygotes in locus |  |  |  |  |  |  |
| --- | --- | --- | --- | --- | --- | --- | --- | --- |
|  |  | 17 | 36 | 23 | 9 | 12 | 67 | average |
| Crete | 30 | 0.3333 | 1.0000 | 0.7333 | 1.0000 | 0.0333 | 0.9667 | 0.6778 |
| Cyprus | 64 | 0.1563 | 0.9844 | 0.1563 | 0.5781 | 0.0000 | 0.9688 | 0.4740 |
| Georgia | 11 | 0.7372 | 0.9090 | 0.6363 | 0.7272 | 0.1818 | 0.8181 | 0.6682 |
| Italy | 12 | 0.5833 | 0.8333 | 0.2500 | 0.7500 | 0.0000 | 0.5000 | 0.4861 |
| Tajikistan | 9 | 0.3333 | 1.0000 | 1.0000 | 0.7778 | 0.0000 | 1.0000 | 0.6852 |
| Other | 6 | 0.5000 | 1.0000 | 0.3333 | 1.0000 | 0.0000 | 0.6667 | 0.5833 |
| All | 136 | 0.3235 | 0.9706 | 0.4118 | 0.7279 | 0.0441 | 0.8971 | 0.5625 |

Heterozygosity was increased in some loci, but decreased in others. Observed heterozygosity was significantly higher than expected for loci 36 and 9 in Crete and 36 and 67 in Cyprus, but significantly lower for loci 17, 23 and 12 in Crete and 17 in Cyprus (Table 2). For other loci, there was no significant difference between observed and expected heterozygosity.

**Table:** Comparison between expected and observed proportion of heterozygotes. HetEXP = expected proportion of heterozygotes, HetOBS = observed proportion of heterozygotes, p(excess) – p-value of heterozygote excess test, p(deficit) – p-value of heterozygote deficit test. All calculation performed in software Genepop. Bold distinguishes significant values. Locus 12 in Cyprus population had only one allele, therefore excess or deficit of heterozygotes was not possible to calculate.

| locus | Crete(N=26) |  |  |  | Cyprus (N=40) |  |  |  |
| --- | --- | --- | --- | --- | --- | --- | --- | --- |
|  | HetEXP | HetOBS | p(excess) | p(deficit) | HetEXP | HetOBS | p(excess) | p(deficit) |
| 17 | 0.6054 | 0.2692 | 0.9974 | **0.0027** | 0.5888 | 0.1750 | 1.0000 | **0.0000** |
| 36 | 0.8150 | 1.0000 | **0.0048** | 1.0000 | 0.5513 | 0.9750 | **0.0000** | 1.0000 |
| 23 | 0.6958 | 0.6923 | 0.9824 | **0.0176** | 0.1420 | 0.1500 | 0.8142 | 1.0000 |
| 9 | 0.6266 | 1.0000 | **0.0000** | 1.0000 | 0.6363 | 0.6250 | 0.4081 | 0.5997 |
| 12 | 0.1477 | 0.0000 | 1.0000 | **0.0007** | 0.0000 | 0.0000 | NA | NA |
| 67 | 0.7177 | 0.9615 | 0.6578 | 0.3422 | 0.6363 | 0.9500 | **0.0000** | 1.0000 |

### Diversity of genotypes

In Tajikistan, the most common genotype had a frequency of 44.44% (4/9), The second most common genotype had frequency of 33.33% (3/9). Other two genotypes had a frequency of 11.11% (1/9). In Italy, the most common genotype had frequency of 16.66%. The other 10 genotypes were found each in one individual. In Georgia, the most common genotype had a frequency of 27% (3/11). The other 8 genotypes were detected only once. In Cyprus we found two genotypes with a frequency of 15.625% (10/64). The third most common genotype was found with a frequency of 12.5% (8/64). The other seven genotypes had a frequency of between 3.13-4.69%. The other 18 genotypes were detected only once (frequency 1.57%). In Crete, the most common genotype had a frequency of 30% (9/30). The second most common genotypes had a frequency of 10% (3/10). There were also found genotypes with a frequency of 6.66% (2/30). We also found 14 genotypes which were found only once.

## Relatedness

We found that almost all offspring genotypes were identical to mother’s (97.87%, 184/188, out of 59 nests). The same genotype in mother and offspring was found in all nests in Albania (2 nests, 10 offspring), Crete (6 nests, 23 offspring), and Italy (10 nests, 38 offspring). In Cyprus, we found one offspring with a different genotype than its mother (out of 89 offspring and 32 nests) and in Tajikistan we found 3 such offspring (out of 9 nests and 28 offspring). However, all offspring for which we detected different genotype than of mother had much lower detection peak for multiple microsatellite loci than most of the analyzed individuals. All four individuals contained at least one allele which is not shared with their mother. Two individuals from Tajikistan both had alleles different than mother in at least one locus. In one individual from Cyprus and two from Tajikistan had a unique allele, which was not found in any other genotyped individual. This suggests that apparent differences between offspring and maternal genotypes were the result of genotyping errors.

## Discussion

*Ceratina dallatoreana* was considered to reproduce parthenogenetically (Daly, 1966, 1983), however, genetic evidence was lacking. We observed parthenogeny in several locations in Mediterranean (Albania, Italy, Crete, Cyprus) and central Asia (Tajikistan), providing evidence for parthenogenesis from a large part of the original geographic range of the species. As males were extremely rare in North Africa (Daly, 1983) and also California, where the species is introduced (Daly, 1966) we can suppose that parthenogeny is prevailing or only mode of reproduction in the whole distributional area.

Thelytokous parthenogeny is generally rare in bees and well-documented only in *Apis mellifera capensis* (Goudie & Oldroyd, 2014; Rabeling & Kronauer, 2013). Outside *Apis*, parthenogeny has been documented only in the *Ceratina* genus of small carpenter bees. This includes evidence for parthenogeny in *C. acantha* ((Slobodchikoff & Daly, 1971), *C. dentipes* (Shell & Rehan, 2019; Snelling, 2003, Mikát unpublished data), *C. parvula* (Terzo et al., 2007, Mikát, unpublished data) and *C. dallatoreana* ((Daly, 1966), this study). These species are not closely related, belonging to different subgenera (Ascher & Pickering, 2020; Rehan & Schwarz, 2015). Species which are considered to be the most closely related to *C. dallatoreana*, such as *C. dentiventris* and *C. sakagamii* do not have skewed sex ratio (Daly, 1983; Terzo, 1998), therefore parthenogeny is probably not the prevailing mode of reproduction in these two species. However for the species *C. rasmonti*, known from only few individuals and closely related to *C. dallatoreana*, males are unknown (Terzo, 1998) therefore this species can be also parthenogenetic, but larger sample size is necessary for proper evaluation. The distribution of parthenogeny in different *Ceratina* lineages suggests that there is a trend for parthenogenesis to arise in the *Ceratina* genus, but future research including sampling of more species and high-resolution phylogeny is necessary for evaluation of frequency and evolution of parthenogenesis in *Ceratina*.

Although we found only few offspring with genotypes that were not identical to genotypes of mothers, we suspect that these cases were the result of genotyping errors. Situations in which offspring had different genotypes than mother were usually not compatible with scenario of sexual reproduction. These results were also incompatible with any mode of parthenogenesis, because we detected alleles in offspring which are not detected in mother. In case of parthenogeny we can suppose allele loss, but not an allele arise.

Offspring resulting from parthenogenesis should bear only alleles which also bear their mother. However, cytology of parthenogeny determines the rate of loss of heterozygosity from mother to offspring (Pearcy et al., 2006). We did not observe any heterozygosity loss – all offspring were genetically identical to their mother when four improperly genotyped individuals were excluded. This strongly supports that offspring are identical clones of their mother. This situation is compatible with two cytological types of parthenogeny: apomixis or automixis with central fusion. Our results are fully compatible with possibility of apomixis. Automixis with central fusion is less probable, as under this scenario, there should be at least some heterozygosity loss due to recombination (Engelstädter, 2017; Goudie & Oldroyd, 2014). Therefore, automixis with central fusion is possible in *C. dallatoreana* only if recombination is missing, its rate is extremely low or if all six our microsatellites are very close to the centromere. Empirical studies on organisms with central fusion automixis using microsatellites showed at least some heterozygosity loss (Fougeyrollas et al., 2015; Rey et al., 2011). Studies of *Apis mellifera capensis* show that homozygotes arise due to recombination, but they often die during early developmental stages – therefore, high heterozygosity is preserved by selection (Goudie & Oldroyd, 2014). As we did not find any case of a homozygote offspring with a heterozygote mother (even in offspring egg stage), apomixis is the more probable mechanism.

We showed that thelytokous parthenogenesis is the prevailing mode of reproduction in *C. dallatoreana*. However, there remains a question as to whether sexual reproduction is only extremely rare, or if it does not occur at all. Existence of males is rarely reported for this species, however, most of reports of males could have been confused with closely related species (Daly, 1983; Terzo, 1998). But males are undoubtedly reported from California, where *C. dallatoreana* is invasive and no similar species are present (Daly, 1966). However, the existence of males alone does not prove their involvement in reproduction. Strictly apomictic species have usually only one or a few genotypes in one location or region (Lorenzo-Carballa & Cordero-Rivera, 2009; Ryskov et al., 2017). However, although we detected some genotypes repeatedly, there was generally high genotype diversity in each location, suggesting that sexual reproduction sometimes occurs in *C. dallatoreana*, although it is likely very rare.

The best documented examples of thelytoky in aculeate Hymenoptera are found among advanced eusocial species, and features of thelytoky are influenced by their social organization (Goudie & Oldroyd, 2018). On the other hand, *Ceratina* bees are mostly facultatively social (Groom & Rehan, 2018; Rehan, 2020). Although most studied species are able to establish social colonies, the larger proportion of the population is solitary and social colonies contain only two or a few females (Groom & Rehan, 2018; Mikát et al., 2022; Rehan et al., 2009; Sakagami & Maeta, 1977). Reversion to strict solitarity also exist in some species (Groom & Rehan, 2018; Mikát et al., 2020). Social nests have not been documented in *C. dallatoreana* to date (Daly, 1966; Mikát et al., 2022), thought the number of nests so far analyzed does not preclude the possibility of very rare sociality in this species. Regardless, it is clear that social nesting is at least very uncommon in this species. This is quite surprising, because parthenogeny should promote social behavior, due to high relatedness between mother and offspring (Hamilton, 1964). Moreover, two other species of *Ceratina* where parthenogeny probably occurs (*C. dentipes* and *C. parvula*) are facultatively social (Mikát et al., 2022; Rehan et al., 2009; Terzo et al., 2007). In last parthenogenetic species, *Ceratina acantha* (Slobodchikoff & Daly, 1971), social status was not examined to this date.

*Ceratina* bees are generally an excellent group for study of social evolution, due to their extensive within and between species variability in social behavior (Groom & Rehan, 2018; Rehan, 2020). Existence of parthenogeny in many *Ceratina* species which are not closely related provides us with further examples of between-species variability in relatedness, a highly important and probably key parameter influencing social evolution (Foster et al., 2006; Hamilton, 1964; Nonacs, 2017). However, better understating of within-colony relatedness and natural history of more species is necessary for understanding co-evolution of thelytoky and sociality.

## Supporting information

supplementary_mikat_dallatoreana

## Acknowledgments

We are grateful to Daniel Benda, Karolína Dobešová, Klára Daňková, Slavomír Dobrotka, Zuzana Dobrotková, Karolína Fazekašová, Tereza Fraňková, Jiří Houska, Jiří Janoušek, Lukáš Janošík, Celie Korittová, Tereza Maxerová, Miroslav Mikát, Blanka Mikátová, Jindra Mrozek, Daniela Reiterová, Tadeáš Ryšan, Vít Procházka, Vojtěch Waldhauser and Jitka Waldhauserová to their assistance in field. We are also grateful to Vít Bureš and Celie Korritová for collecting additional bees. We are grateful to Jesse Huisken for feedback to manuscript. The Grant Agency of Charles University (Grant GAUK 764119/2019) and the Specific University Research Project Integrative Animal Biology (Grant SVV 260571/2021) supported this research.

## References

Alavi, Y., Rooyen, A. van, Elgar, M. A., Jones, T. M., & Weeks, A. R. (2018). Novel microsatellite markers suggest the mechanism of parthenogenesis in Extatosoma tiaratum is automixis with terminal fusion. Insect Science, 25(1), 24–32. https://doi.org/10.1111/1744-7917.12373

Ascher, John. S., & Pickering, J. (2020). Discover Life bee species guide and world checklist (Hymenoptera: Apoidea: Anthophila). http://www.discoverlife.org/mp/20q?guide=Apoidea_species

Daly, H. V. (1966). Biological studies on Ceratina dallatorreana, an alien bee in California which reproduces by parthenogenesis (Hymenoptera: Apoidea). Annals of the Entomological Society of America, 59(6), 1138–1154.

Daly, H. V. (1983). Taxonomy and ecology of Ceratinini of North Africa and the Iberian Peninsula (Hymenoptera: Apoidea). Systematic Entomology, 8(1), 29–62. https://doi.org/10.1111/j.1365-3113.1983.tb00466.x

Engelstädter, J. (2008). Constraints on the evolution of asexual reproduction. BioEssays, 30(11–12), 1138–1150. https://doi.org/10.1002/bies.20833

Engelstädter, J. (2017). Asexual but Not Clonal: Evolutionary Processes in Automictic Populations. Genetics, 206(2), 993–1009. https://doi.org/10.1534/genetics.116.196873

Foster, K. R., Wenseleers, T., Ratnieks, F. L., & Queller, D. C. (2006). There is nothing wrong with inclusive fitness. Trends in Ecology & Evolution, 21(11), 599–600.

Fougeyrollas, R., Dolejšová, K., Sillam-Dussès, D., Roy, V., Poteaux, C., Hanus, R., & Roisin, Y. (2015). Asexual queen succession in the higher termite Embiratermes neotenicus. Proceedings of the Royal Society B: Biological Sciences, 282(1809), 20150260. https://doi.org/10.1098/rspb.2015.0260

Fujita, M. K., Singhal, S., Brunes, T. O., & Maldonado, J. A. (2020). Evolutionary Dynamics and Consequences of Parthenogenesis in Vertebrates. Annual Review of Ecology, Evolution, and Systematics, 51(1), 191–214. https://doi.org/10.1146/annurev-ecolsys-011720-114900

Gerber, H. S., & Klostermeyer, E. C. (1970). Sex control by bees: A voluntary act of egg fertilization during oviposition. Science, 167(3914), 82–84.

Gokhman, V. E., & Kuznetsova, V. G. (2018). Parthenogenesis in Hexapoda: Holometabolous insects. Journal of Zoological Systematics and Evolutionary Research, 56(1), 23–34.

Goudie, F., & Oldroyd, B. P. (2014). Thelytoky in the honey bee. Apidologie, 45(3), 306–326. https://doi.org/10.1007/s13592-013-0261-2

Goudie, F., & Oldroyd, B. P. (2018). The distribution of thelytoky, arrhenotoky and androgenesis among castes in the eusocial Hymenoptera. Insectes Sociaux, 65(1), 5–16.

Groom, S. V. C., & Rehan, S. M. (2018). Climate-mediated behavioural variability in facultatively social bees. Biological Journal of the Linnean Society, 125(1), 165–170. https://doi.org/10.1093/biolinnean/bly101

Hamilton, W. D. (1964). The genetical evolution of social behaviour. I. Journal of Theoretical Biology, 7(1), 1–16. https://doi.org/10.1016/0022-5193(64)90038-4

Heinze, J. (2008). The demise of the standard ant (Hymenoptera: Formicidae). Myrmecological News, 11, 9–20.

Hörandl, E., Bast, J., Brandt, A., Scheu, S., Bleidorn, C., Cordellier, M., Nowrousian, M., Begerow, D., Sturm, A., & Verhoeven, K. (2020). Genome Evolution of Asexual Organisms and the Paradox of Sex in Eukaryotes. Evolutionary Biology—A Transdisciplinary Approach, 133–167.

Kooi, C. J. van der, Matthey-Doret, C., & Schwander, T. (2017). Evolution and comparative ecology of parthenogenesis in haplodiploid arthropods. Evolution Letters, 1(6), 304–316. https://doi.org/10.1002/evl3.30

Liegeois, M., Sartori, M., & Schwander, T. (2021). Extremely Widespread Parthenogenesis and a Trade-Off Between Alternative Forms of Reproduction in Mayflies (Ephemeroptera). Journal of Heredity, 112(1), 45–57. https://doi.org/10.1093/jhered/esaa027

Lorenzo-Carballa, M. O., & Cordero-Rivera, A. (2009). Thelytokous parthenogenesis in the damselfly Ischnura hastata (Odonata, Coenagrionidae): Genetic mechanisms and lack of bacterial infection. Heredity, 103(5), 377–384. https://doi.org/10.1038/hdy.2009.65

Michener, C. D. (1960). Notes on the Biology and Supposed Parthenogenesis of Halictine Bees from the Australian Region. Journal of the Kansas Entomological Society, 33(2), 85–96. JSTOR.

Mikát, M., Benda, D., Korittová, C., Mrozková, J., Reiterová, D., Waldhauserová, J., Brož, V., & Straka, J. (2020). Natural history and maternal investment of Ceratina cucurbitina, the most common European small carpenter bee, in different European regions. Journal of Apicultural Research, 1–12.

Mikát, M., Fraňková, T., Benda, D., & Straka, J. (2022). Evidence of sociality in European small Carpenter bees (Ceratina). Apidologie, 53(2), 18. https://doi.org/10.1007/s13592-022-00931-8

Mikát, M., Janošík, L., Černá, K., Matoušková, E., Hadrava, J., Bureš, V., & Straka, J. (2019). Polyandrous bee provides extended offspring care biparentally as an alternative to monandry based eusociality. Proceedings of the National Academy of Sciences, 116(13), 6238–6243. https://doi.org/10.1073/pnas.1810092116

Mikát, M., Matoušková, E., & Straka, J. (2021). Nesting of Ceratina nigrolabiata, a biparental bee. Scientific Reports, 11(1), 5026. https://doi.org/10.1038/s41598-021-83940-4

Neiman, M., Meirmans, S., & Meirmans, P. (2009). What can asexual lineage age tell us about the maintenance of sex? Annals of the New York Academy of Sciences, 1168(1), 185–200.

Neiman, M., & Schwander, T. (2011). Using Parthenogenetic Lineages to Identify Advantages of Sex. Evolutionary Biology, 38(2), 115–123. https://doi.org/10.1007/s11692-011-9113-z

Nonacs, P. (2017). Go High or Go Low? Adaptive Evolution of High and Low Relatedness Societies in Social Hymenoptera. Frontiers in Ecology and Evolution, 5. https://doi.org/10.3389/fevo.2017.00087

Normark, B. B. (2003). The evolution of alternative genetic systems in insects. Annual Review of Entomology, 48(1), 397–423. https://doi.org/10.1146/annurev.ento.48.091801.112703

Pearcy, M., Hardy, O., & Aron, S. (2006). Thelytokous parthenogenesis and its consequences on inbreeding in an ant. Heredity, 96(5), 377–382. https://doi.org/10.1038/sj.hdy.6800813

Rabeling, C., & Kronauer, D. J. (2013). Thelytokous parthenogenesis in eusocial Hymenoptera. Annual Review of Entomology, 58, 273–292.

Rehan, S. M. (2020). Small Carpenter Bees (Ceratina). In C. K. Starr (Ed.), Encyclopedia of Social Insects (pp. 1–4). Springer International Publishing. https://doi.org/10.1007/978-3-319-90306-4_106-1

Rehan, S. M., & Richards, M. H. (2010). Nesting biology and subsociality in Ceratina calcarata (Hymenoptera: Apidae). The Canadian Entomologist, 142(1), 65–74. https://doi.org/10.4039/n09-056

Rehan, S. M., Richards, M. H., & Schwarz, M. P. (2009). Evidence of Social Nesting in the Ceratina of Borneo (Hymenoptera: Apidae). Journal of the Kansas Entomological Society, 82(2), 194–209. https://doi.org/10.2317/JKES809.22.1

Rehan, S., & Schwarz, M. (2015). A few steps forward and no steps back: Long-distance dispersal patterns in small carpenter bees suggest major barriers to back-dispersal. Journal of Biogeography, 42(3), 485–494. https://doi.org/10.1111/jbi.12439

Rey, O., Loiseau, A., Facon, B., Foucaud, J., Orivel, J., Cornuet, J.-M., Robert, S., Dobigny, G., Delabie, J. H. C., Mariano, C. D. S. F., & Estoup, A. (2011). Meiotic Recombination Dramatically Decreased in Thelytokous Queens of the Little Fire Ant and Their Sexually Produced Workers. Molecular Biology and Evolution, 28(9), 2591–2601. https://doi.org/10.1093/molbev/msr082

Rousset, F. (2020). Genepop Version 4.7. 0. Institut des Sciences de l’Evolution de Montpellier, Université de ….

Ryskov, A. P., Osipov, F. A., Omelchenko, A. V., Semyenova, S. K., Girnyk, A. E., Korchagin, V. I., Vergun, A. A., & Murphy, R. W. (2017). The origin of multiple clones in the parthenogenetic lizard species Darevskia rostombekowi. PLOS ONE, 12(9), e0185161. https://doi.org/10.1371/journal.pone.0185161

Sakagami, S. F., & Maeta, Y. (1977). Some presumably presocial habits of Japanese Ceratina bees, with notes on various social types in Hymenoptera. Insectes Sociaux, 24(4), 319–343. https://doi.org/10.1007/BF02223784

Schwander, T., & Crespi, B. J. (2009). Multiple Direct Transitions from Sexual Reproduction to Apomictic Parthenogenesis in Timema Stick Insects. Evolution, 63(1), 84–103. https://doi.org/10.1111/j.1558-5646.2008.00524.x

Shell, W. A., & Rehan, S. M. (2019). Invasive range expansion of the small carpenter bee, Ceratina dentipes (Hymenoptera: Apidae) into Hawaii with implications for native endangered species displacement. Biological Invasions, 21(4), 1155–1166. https://doi.org/10.1007/s10530-018-1892-z

Shukla, S., Shilpa, M. C., & Gadagkar, R. (2013). Virgin wasps develop ovaries on par with mated females, but lay fewer eggs. Insectes Sociaux, 60(3), 345–350.

Slobodchikoff, C. N., & Daly, H. V. (1971). Systematic and evolutionary implications of parthenogenesis in the Hymenoptera. American Zoologist, 11(2), 273–282.

Snelling, R. R. (2003). Bees of the Hawaiian Islands, exclusive of Hylaeus (Nesoprosopis)(Hymenoptera: Apoidea). Journal of the Kansas Entomological Society, 342–356.

Stenberg, P., & Saura, A. (2009). Cytology of asexual animals. In Lost sex (pp. 63–74). Springer.

Stubblefield, J. W., & Seger, J. (1994). Sexual dimorphism in the Hymenoptera. In The differences between the sexes. (pp. 71–103). Cambridge University Press, Cambridge.

Terzo, M. (1998). Annotated list of the species of the genus Ceratina (Latreille) occuring in the Near East, with descriptions of new species (Hymenoptera: Apoidea: Xylocopinae). Linzer Biologische Beiträge, 30(2), 719–743.

Terzo, M., Iserbyt, S., & Rasmont, P. (2007). Révision des Xylocopinae (Hymenoptera: Apidae) de France et de Belgique. Annales de La Société Entomologique de France, 43, 445–491. http://www.tandfonline.com/doi/abs/10.1080/00379271.2007.10697537

Terzo, M., & Rasmont, P. (2004). Biogéographie et systématique des abeilles rubicoles du genre Ceratina Latreille au Turkestan (Hymenoptera, Apoidea, Xylocopinae). Annales de La Société Entomologique de France (N.S.), 40(2), 109–130. https://doi.org/10.1080/00379271.2004.10697410

Terzo, M., & Rasmont, P. (2011). Atlas of the European Bees: Genus Ceratina. Atlas Hymenoptera - Atlas of the European Bees - STEP Project. http://www.atlashymenoptera.net/page.asp?id=192

Thierry, L. (2013). Adaptive significance and long-term survival of asexual lineages 2. Evolutionary Biology, 40(3), 450–460.

Tsutsui, Y., Maeto, K., Hamaguchi, K., Isaki, Y., Takami, Y., Naito, T., & Miura, K. (2014). Apomictic parthenogenesis in a parasitoid wasp Meteorus pulchricornis, uncommon in the haplodiploid order Hymenoptera. Bulletin of Entomological Research, 104(3), 307.

Tvedte, E. S., Logsdon, J. M., & Forbes, A. A. (2019). Sex loss in insects: Causes of asexuality and consequences for genomes. Current Opinion in Insect Science, 31, 77–83. https://doi.org/10.1016/j.cois.2018.11.007

Vavre, F., de Jong, J. H., & Stouthamer, R. (2004). Cytogenetic mechanism and genetic consequences of thelytoky in the wasp Trichogramma cacoeciae. Heredity, 93(6), 592–596. https://doi.org/10.1038/sj.hdy.6800565

Vorburger, C. (2014). Thelytoky and sex determination in the hymenoptera: Mutual constraints. Sexual Development, 8(1–3), 50–58.

Wenseleers, T., & Van Oystaeyen, A. (2011). Unusual modes of reproduction in social insects: Shedding light on the evolutionary paradox of sex. BioEssays, 33(12), 927–937. https://doi.org/10.1002/bies.201100096

