## supplementary_mikat_dallatoreana for "Genetic evidence for parthenogenesis in small carpenter bee, *Ceratina dallatoreana* in its native distribution area"

**Faunistic notes:**

**Tajikistan:** Terzo and Rasmont (2004) do not list *C. dallatoreana* species from Tajikistan. We found this species commonly in several locations in Tajikistan: Dushambe (38.4568586N, 68.7844932E, Dzarteppa (38.0197706N, 69.3592158E), Rozien (38.2706964N, 69.2591528E), and Sharison (39.7800442N, 68.8118569E). On the other hand, we did not find *C. dallatoreana* in Varzob (38.8120675N, 68.8228925E) also several locations around Anzob (39.1180006N, 68.8483583E, 39.1589481N, 68.8421356E, 39.1925336N, 68.6638725E), although there other *Ceratina* species were present. Generally, *C. dallatoreana* was present in low and middle elevations (up to around 1500 m) and absent in Zarafsah range. We can suppose that *C. dallatoreana* is common in most of Tajikistan outside high mountains.

**Georgia**: Terzo and Rasmont (2011) show only one data point from Albania. We found *C. dallatoreana* in several locations. Although this species is much less commons than *C. cucurbitina* in this country (Mikát et al., 2020), we suppose that it widespread across whole country.

**Georgia**: Only one location according to map (Terzo and Rasmont, 2011). However, We found it in several locations across the country. This species was dominant *Ceratina* species in the most south-east edge of country (Vashlovani).

**Microsatellite primers:**

**Table S1.** Features of microsatellite loci tested on *C. dallatoreana*. These microsatellite primers were primary developed on C. nigrolabiata (Mikát et al., 2019)

| Locus  N | Forward primer | Reverse primer | Motive of repetition | Approx. number of repetitions | Results on *C. dallatoreana* |
| --- | --- | --- | --- | --- | --- |
| 17 | 6FAM-CGATTGCCTAGTCTCCTCCA | GTTTGGTGGTACGCGGAATCTAAT | tc | 11 | Used |
| 67 | PET-CGTGACGCGACTTTCATTC | GTTTCCTCTCACCCACTCTTACTGTCA | ag | 10+5 | Used |
| 23 | VIC-GCACTCGTTCCTCCCTTTC | GTTTGAACACTGGTGCACCTCAGA | tc | 10 | Used |
| 28 | NED-CTCGAACTAATTACCCGACGA | GTTTTCCTACATCCCTGTCAGC | ag | 10 | Not polymorphic |
| 30 | 6FAM-TAGAATCCTTGGAAGATTAGTGGT | GTTTACCCCGGGTCCCTAATTC | ag | 10 | Polymorphic, but overlap with other primers with same color |
| 27 | PET-TTTCCCTGAAGCAAATGCAG | GTTTAACTCCGTACGGCCTTCAG | tc | 10 | Polymorphic, but low quality of amplification |
| 9 | VIC-TATGCGATTAGCTAGCGCGG | GTTTCCCTGGCCGTACTCTTATGC | ag | 17 | Used |
| 12 | VIC-AGGATGGACCGGACGGAATA | GTTTGATGTTCCCTGGGTCAGGTG | ct | 13 | Used |
| 43 | NED-AACCGGATTCCTACTACGGG | GTTTCTTCGTGATGGAAATCGTGA | gac | 9 | Not Polymorphic |
| 36 | 6FAM-CCTTCTGCCCTACCTTGAAA | GTTTCAGAACCAGAGGGAGACGAA | ag | 10 | Used |
| 7 | PET-TTGGCGAGATTATGGAAACG | GTTTCAGTTTTATATCTCGTTGCCTTTT | aatg | 11 | Not Polymorphic |
| 8 | VIC-ACGTGTACACCGACTACGTG | GTTTAATTCTGGCCAGGTTGGAG | tc | 11 | Polymorphic, but interaction with other primers |
| 46 | NED-GCGAACGTATCGTTCTCACA | GTTTAACGTTAACCACACCTTAAACG | agcg/ag | 17 | Not successful amplification |
| 51 | PET-CACGCTGTCCCTACGAGTTT | GTTTCCCGGTGAACGTCCAAAA | tc | 13 | Polymorphic, but low quality of amplification |

Table S2: Features of successfully amplified microsatellites for *C. dallatoreana*

| Locus N | N Alleles | Range | Overall heterozygosity |
| --- | --- | --- | --- |
| 17 | 16 | 120-162 | 0.3235 |
| 36 | 18 | 225-279 | 0.9706 |
| 23 | 5 | 97-105 | 0.4118 |
| 9 | 12 | 208-256 | 0.7279 |
| 12 | 3 | 207-214 | 0.0441 |
| 67 | 6 | 95-105 | 0.8971 |

**References**

Mikát, M., Benda, D., Korittová, C., Mrozková, J., Reiterová, D., Waldhauserová, J., Brož, V., Straka, J., 2020. Natural history and maternal investment of Ceratina cucurbitina, the most common European small carpenter bee, in different European regions. J. Apic. Res. 1–12.

Mikát, M., Janošík, L., Černá, K., Matoušková, E., Hadrava, J., Bureš, V., Straka, J., 2019. Polyandrous bee provides extended offspring care biparentally as an alternative to monandry based eusociality. Proc. Natl. Acad. Sci. 116, 6238–6243. https://doi.org/10.1073/pnas.1810092116

Terzo, M., Rasmont, P., 2011. Atlas of the European Bees: genus Ceratina [WWW Document]. Atlas Hymenopt. - Atlas Eur. Bees - STEP Proj. URL http://www.atlashymenoptera.net/page.asp?id=192 (accessed 2.25.19).

Terzo, M., Rasmont, P., 2004. Biogéographie et systématique des abeilles rubicoles du genre Ceratina Latreille au Turkestan (Hymenoptera, Apoidea, Xylocopinae). Ann. Société Entomol. Fr. NS 40, 109–130. https://doi.org/10.1080/00379271.2004.10697410
